# A potent MAPK13-14 inhibitor prevents airway inflammation and mucus production

**DOI:** 10.1101/2023.05.26.542451

**Authors:** Shamus P. Keeler, Kangyun Wu, Yong Zhang, Dailing Mao, Ming Li, Courtney A. Iberg, Stephen R. Austin, Samuel A. Glaser, Jennifer Yantis, Stephanie Podgorny, Steven L. Brody, Joshua R. Chartock, Zhenfu Han, Derek E. Byers, Arthur G. Romero, Michael J. Holtzman

**Affiliations:** Pulmonary and Critical Care Medicine, Department of Medicine, Washington University School of Medicine, St. Louis, MO 63110; Department of Cell Biology and Physiology, Washington University School of Medicine, St. Louis, MO 63110; NuPeak Therapeutics Inc., St. Louis, MO 63105

**Keywords:** asthma, chronic obstructive pulmonary disease (COPD), drug discovery, minipig model, respiratory viral infection

## Abstract

Common respiratory diseases continue to represent a major public health problem, and much of the morbidity and mortality is due to airway inflammation and mucus production. Previous studies indicated a role for mitogen-activated protein kinase 14 (MAPK14) in this type of disease, but clinical trials are unsuccessful to date. Our previous work identified a related but distinct kinase known as MAPK13 that is activated in respiratory airway diseases and is required for mucus production in human cell-culture models. Support for MAPK13 function in these models came from effectiveness of *MAPK13* versus *MAPK14* gene-knockdown and from first-generation MAPK13-14 inhibitors. However, these first-generation inhibitors were incompletely optimized for blocking activity and were untested in vivo. Here we report the next generation and selection of a potent MAPK13-14 inhibitor (designated NuP-3) that more effectively down-regulates type-2 cytokine-stimulated mucus production in air-liquid interface and organoid cultures of human airway epithelial cells. We also show that NuP-3 treatment prevents respiratory airway inflammation and mucus production in new minipig models of airway disease triggered by type-2 cytokine challenge or respiratory viral infection. The results thereby provide the next advance in developing a small-molecule kinase inhibitor to address key features of respiratory disease.

**New and noteworthy:** This study describes the discovery of a potent MAPK13-14 inhibitor and its effectiveness in models of respiratory airway disease. The findings thereby provide a scheme for pathogenesis and therapy of lung diseases (e.g., asthma, COPD, Covid-19, post-viral and allergic respiratory disease) and related conditions that implicate MAPK13-14 function. The findings also refine a hypothesis for epithelial and immune cell functions in respiratory disease that features MAPK13 as a possible component of this disease process.

## INTRODUCTION

Respiratory diseases, commonly in the form of asthma and COPD, remain leading causes of morbidity and mortality in the U.S. and worldwide despite current therapeutic approaches (1, 2). Moreover, there is growing recognition that these diseases are linked to obstruction of the airways with inflammatory cells and mucus (3-9). Therefore, attenuation of airway inflammation and mucus production would likely improve airflow, muco-ciliary clearance, and host defense (10, 11) and thereby meet a target endpoint of decreasing exacerbation and progression in airway diseases. However, these concepts cannot be fully validated until there are specific and effective therapies to normalize airway inflammation and associated mucus production. In that context, we used human epithelial cell models and kinase arrays to identify a stress kinase known as mitogen-activated protein kinase 13 (MAPK13) as a requirement for basal-epithelial stem cell (basal-ESC) transition to mucous cells during type-2 immune stimulation in cell models and patients with COPD. (12, 13). These findings stand in some contrast to the role of closely-related MAPK14 as part of the immune response that is conventionally linked to inflammatory phenotypes, including respiratory inflammation (14-18) and mucus production (19-24). However, even highly potent MAPK14 inhibitors when given alone have been ineffective in clinical trials of COPD patients (25).

Together, this work suggested a therapeutic advantage of a combined MAPK13-14 inhibitor for treatment of respiratory disease. However, to our knowledge, no potent MAPK13-14 inhibitors were yet available (26, 27). Here we report the use of structure-based drug design to extend our previous work on first-generation compounds and arrive at a second-generation MAPK13-14 inhibitor designated NuP-3. We demonstrate that NuP-3 exhibits more favorable target-binding and blockade in enzyme assays, potent inhibition of mucus production in human airway epithelial cell culture, and effective attenuation of airway inflammation and mucus production in type-2 cytokine-challenge and viral-infection models of airway disease in minipigs in vivo.

## MATERIALS AND METHODS

### MAPK inhibitor generation and assay

MAPK inhibitors were developed as a chemical analog series using structure-based drug design as introduced previously (12). The entire series was subjected to a screening funnel that began with assessments of chemical properties and MAPK13 and MAPK14 enzyme inhibition assays. Enzyme inhibition assays were performed using the HotSpot assay platform as described previously (28). Compounds were reconstituted in DMSO vehicle (at least 1:1000 vol/vol) for cell-culture model experiments and were dissolved in Ensure® nutritional supplement for minipig model experiments. For each preparation, compound purity was verified using LC-MS and NMR.

### Epithelial cell culture

Human tracheal and bronchial epithelial cells (hTECs) were isolated by enzymatic digestion, seeded onto permeable filter supports, and grown as described previously (12, 13). For the present experiments, cells were cultured in 24-well Transwell plates (6.5-mm diameter inserts, 0.4 µm pore size) from Corning (Corning, NY) with 2% NuSerum medium (29) supplemented with Primocin (50 µg/ml, (InvivoGen, San Diego, CA), and retinoic acid (1 x 10^−8^ M, Sigma, St. Louis, MO) with or without human IL-13 (50 ng/ml, Peprotech, Rocky Hill, NJ) under submerged conditions for 7 d and then air-liquid interface conditions for 21 d. Cells were cultured in the presence of test compound or vehicle that was added 2 d before addition of IL-13 and was re-added with each medium change/IL-13 treatment (twice per week). For each condition, the numbers of viable cells were monitored using a resazurin-based CellTiter-Blue Cell Viability Assay (#G8080, Promega, Madison, WI). In addition, hTECs were cultured under 3D-Matrigel conditions to permit organoid formation as described previously (13). In these experiments, the IL-13 concentration was set at 1 ng/ml that was optimal for well-differentiated organoid formation and preservation. Test compound or vehicle was added as described above for air-liquid interface conditions. Compound effect on mucus production was based on target protein and mRNA levels that were determined as described below.

### ELISA

To assess mucus production at the protein level, the levels of mucin 5AC (MUC5AC) and chloride channel accessory 1 (CLCA1) were determined in samples that were assayed in triplicate on 96-well white half-area flat-bottom ELISA plates (Greiner Bio-One North America) after incubation at 37 °C overnight. The assay for MUC5AC was performed using biotin-conjugated mouse anti-MUC5AC mAb (45M1, Invitrogen, #MA5-12175) as described previously (12). The assay for CLCA1 was performed using rabbit anti-human CLCA1 (amino acid 33-63) and HRP-conjugated goat anti-rabbit IgG antibody (Santa Cruz Biotechnology) as described previously (12). Assay reactions were developed using the GLO substrate kit (R&D Company) and quantified by comparison to a standard curve with recombinant proteins.

### RNA analysis

To assess mucus production at the RNA level, RNA was purified from cell cultures using the RNeasy mini kit (Qiagen) and corresponding cDNA was generated using the High-Capacity cDNA Archive kit (Life Technologies). We quantified target mRNA levels using real-time qPCR assay with specific fluorogenic probe-primer combinations and Fast Universal PCR Master Mix systems (Applied Biosystems) with human-specific forward and reverse primers and probes as defined in **Supplemental Table 1**. All samples were assayed using the 7300HT or QuantStudio 6 Fast Real-Time PCR System and analyzed using Fast System Software (Applied Biosystems). All real-time PCR data was normalized to the level of *GAPDH* mRNA. Values were expressed as fold-change based on the delta-delta Ct method as described previously (30). For subsequent experiments in minipigs, mRNA targets and viral titer were monitored using the same approach. For those experiments, RNA was purified from homogenized lung tissue using Trizol (Invitrogen) and processed and analyzed as described above, and viral RNA was quantified by copy number using an *SeV-NP*-expressing plasmid as an internal standard. The minipig-specific and virus-specific primers and probes are also defined in **Supplemental Table 1**.

### X-ray crystallography

X-ray crystallography to determine the structure of MAPK-compound co-complexes was performed as described previously (12) with slight modification. In particular, due to the slow association rate of NuP-3 binding to MAPK13, a distinct co-crystallization strategy was applied to prepare a complex crystal. In brief, non-phosphorylated MAPK13 was separated by a MonoQ column as described previously (12) and incubated with NuP-3 at final 1 mM suspension overnight at 4 °C. Then, the complex protein solution was concentrated to 20 mg/ml in 20 mM Hepes, pH 7.4, 150 mM NaCl, and 5% glycerol. Crystals were grown by hanging drop vapor diffusion with the reservoir containing 8∼12 % PEG 4K or 6K, 0.15 M ammonium sulfate and 0.1 M sodium citrate. Diffractable crystals were further optimized by micro-seeding. Data were collected from Advanced Light Source beamline 4.2.2 and processed by XDS/Aimless at 3.2 Å resolution with space group in P2. Final structure was solved by molecular replacement with Rfree/R of 0.324/0.264. The analysis showed four MAPK13–NuP-3 molecules in one asymmetric unit, wherein each active site presented slightly different interactions around its ligand. Molecular graphics figures were produced using PyMOLversion 2.5.2 software.

### Biolayer interferometry (BLI) analysis

For BLI analysis, MAPK-compound binding was performed as described previously (12) using MAPK proteins that were C-terminal biotin tagged. For this approach, codon optimized DNA fragments corresponding to protein sequences of MAPK13 (Uniprot ID, O15264) and MAPK14 (Uniprot ID, Q16539) were synthesized by Integrated DNA Technologies (IDT) and cloned into modified pET28a vectors to introduce the biotin acceptor tag on the C-terminus. Each construct was co-expressed with pBirAcm plasmid (Avidity LLC) in BL21 gold (DE3) *E. coli* cells supplied with 50 µM D-biotin. All biotinylated proteins were purified to >90% purity by chromatography and stored at -80 °C. Kinetic titration assays were performed on the Octet Red384 (ForteBio) system at 25 °C with the running buffer containing 20 mM Hepes, pH 7.4, 140 mM NaCl, 0.02% Tween20, 1 mg/ml BSA, and 2% DMSO. Data were collected in a kinetic titration approach (31, 32) rather than a traditional parallel method. Thus, one Super streptavidin biosensor immobilized with biotinylated MAPK proteins was sequentially dipped into a series of wells having increasing concentration (0.0625 to 1 µM) of compound along with short dissociation after each association step. In addition, double subtraction was applied by using an empty sensor quenched by biocytin and buffer wells dipped by ligand captured biosensor as reference senor and well, respectively. Data were processed with the Octet data analysis 9 package and exported for curve fitting and calculations in the BiaEvaluation software (Biacore; RRID: SCR_015936).

### Pharmacokinetic (PK) analysis and formulation

Compound levels were determined in plasma samples obtained from minipigs (Bama strain) after a single dose at 2 mg/kg and 0.4 mg/ml in Ensure nutritional supplement administered orally by gavage. Bioanalytical compound levels were determined using a Prominence Degasser DGU-20A5T(C) HPLC and an AB Sciex Triple Quad 5500 LC/MS/MS instrument. For NuP-3, limit of detection was 1.1 nM. For these PK experiments and subsequent testing in minipig models in vivo, NuP-3 was prepared and administered as the free base form of the compound. For testing in these models, NuP-3 was administered per os (p.o.) using syringe-feeding to minimize procedural stress.

### Minipig models

Male and female Yucatan minipigs (21-25 kg and 7-8 weeks of age) were obtained from Sinclair Research and housed in environmentally controlled animal care facilities. Minipigs were acclimated for 2 weeks before experiments. Animal husbandry and experimental procedures were approved by the Animal Studies Committees of Washington University School of Medicine in accordance with the guidelines from the National Institutes of Health. Animals were randomly assigned to either treatment or control groups prior to the start of all experiments. Minipigs were anesthetized using tiletamine/zolasepam (Telazol, 4.4 mg/kg), ketamine (2.2 mg/kg), and xylazine (2.2 mg/kg) administered intramuscularly, intubated with a 6.5-7.0-mm ID endotracheal tube, and maintained under 2-3% isoflurane anesthesia. Minipigs were mechanically ventilated using a Drager anesthesia workstation for each challenge procedure. Segmental cytokine challenge was performed on Study Days 0 and 14 using recombinant porcine IL-13 (0.25 mg in 6 ml of PBS) with 2 ml delivered to each of three subsegments (right caudal and accessory lobes) via a fiber-optic bronchoscope (Olympus Model LF-2, 3.8-mm OD). Minipigs were maintained with reverse Trendelenburg position for 30 min to prevent drainage from the right lung before anesthesia was discontinued. The subsequent response to IL-13 challenge was monitored in bronchoalveolar lavage (BAL) fluid and cells at Study Days 2, 4, 7, 14, 16, and 18. Controls included samples from three unchallenged subsegments (left caudal lobes) on Study Day 0 and each of the other Study Day sampling times. BAL was performed by instillation and aspiration of 6-ml aliquots of PBS into left and right anterior lung segments. In some experiments, lungs were removed for tissue analysis on Study Day 16. For treatment, minipigs were dosed orally using syringe-feeding with NuP-3 (2 mg/kg twice per day) in 10 ml of Ensure nutrition shake or with an equivalent amount of Ensure alone on Study Days 12-17. BAL and tissue samples were processed for ELISA and RNA analysis as described above. In addition, BAL samples were analyzed for total and differential cell counts, and tissue samples were processed for immunostaining as described below. The cytokine experiments were structured with sequential challenges in each individual animal to allow the results of drug treatment to be compared to prior nondrug-treated outcomes within the same animal and to results from other strictly control-treated animals. This approach decreased the number of animals used in the study but limited the application of investigator blinding in the experiments. For respiratory virus infection, Sendai virus (SeV) was prepared as described previously (33) and delivered in the same manner as IL-13 via bronchoscopy using 1.3 x 10^7^ pfu in 1 ml of PBS with 1 ml delivered to each of three subsegments on Study Day 0. For analysis, BAL was performed with 6-ml aliquots of PBS in each segment on Study Days 0, 2, 4, 7, 14, and 21, and tissue analysis was obtained on Study Day 7.

### Tissue staining and microscopy

Lung tissue was fixed with 10% formalin, embedded in paraffin, cut into 5-μm thick sections, and adhered to charged slides. Sections were deparaffinized in Fisherbrand CitriSolv (Fisher Scientific), hydrated, and heat-treated with antigen unmasking solution (Vector Laboratories). Immunostaining was performed with MUC5AC mAb (Invitrogen MA5-12175; RRID: AB_10978001), rabbit anti-human CLCA1 antibody (aa 33-63), and rabbit anti-MUC5B antibody (Abcam ab87276; RRID: AB_1951740). Primary antibodies were detected with secondary antibodies labeled with Alexa Fluor 488 (ThermoFisher Scientific) or Alexa Fluor 555 (ThermoFisher Scientific) followed by DAPI counterstaining. Slides were imaged by immunofluorescent microscopy using an Olympus BX51, and staining was quantified using a NanoZoomer S60 slide scanner (Hamamatsu) and ImageJ software version 1.53v (NIH, RRID: SCR_003070, https://imagej.net/ij/index.html) as described previously (34-36).

### Statistical analysis

All data are presented as mean and S.E.M. and are representative of at least three experiments with at least 5 data points per experimental condition. For cell and molecular data, unpaired *t*-test with Bonferroni correction as well as mixed-model repeated measures analysis of variance with Tukey correction for multiple comparisons were used to assess statistical significance between means. In all cases, significance threshold was set at *P* < 0.05.

## RESULTS

### MAPK13 inhibitor generation

To develop a small-molecule inhibitor of MAPK13, we pursued a drug-design strategy to modify a MAPK14 inhibitor parent compound (BIRB-796; NuP-43 in our chemical library) (37) based on NuP-43 chemical structure, NuP-43–MAPK14 co-crystal structure (Protein Data Bank Entry 1KV2), and our predicted binding interactions for NuP-43 to MAPK13 (**Fig. 1a**). In earlier work, we replaced the naphthalene moiety with smaller aromatic rings and halogen substitution to improve fit into the ATP-binding site and decrease reactive metabolite formation and associated toxicity found with the parent compound (38-41). These changes generated first-generation compounds with a modest (3-7-fold) increase in MAPK13 inhibitory activity (12). Here we also modified additional sites that were predicted to interact at the left-hand hinge region key to MAPK specificity (42, 43) and the right-hand allosteric pocket linked to a favorable DFG-out conformation (44) to generate a larger library of chemical analogs (n=428) as recently published (45). These analogs entered a screening funnel that started with a check for physical chemical characteristics based on molecular weight, Lipinski’s Rule of 5, Veber criteria (46), partition coefficient (logP), and topological polar surface area (tPSA) (**Fig. 1b**). Acceptable analogs (n=205) were assessed in MAPK13 and MAPK14 enzyme assays that identified potent inhibitors of MAPK13 and MAPK14 (**Fig. 1c**). Favorable inhibitors from this group (n=28) underwent a cell-based screen for blocking IL-13-stimulated mucus production from human tracheobronchial epithelial cells (hTECs) under air-liquid interface primary-culture conditions. Results showed that mucus inhibition (marked by MUC5AC level) correlated with MAPK13 but not MAPK14 blocking activity (**Fig. 1d**) consistent with our previous results (12). These combined screening tests served as a MAPK13-guided pathway to selection of NuP-3 as a candidate for a combined MAPK13-14 inhibitor.

**Figure 1.**
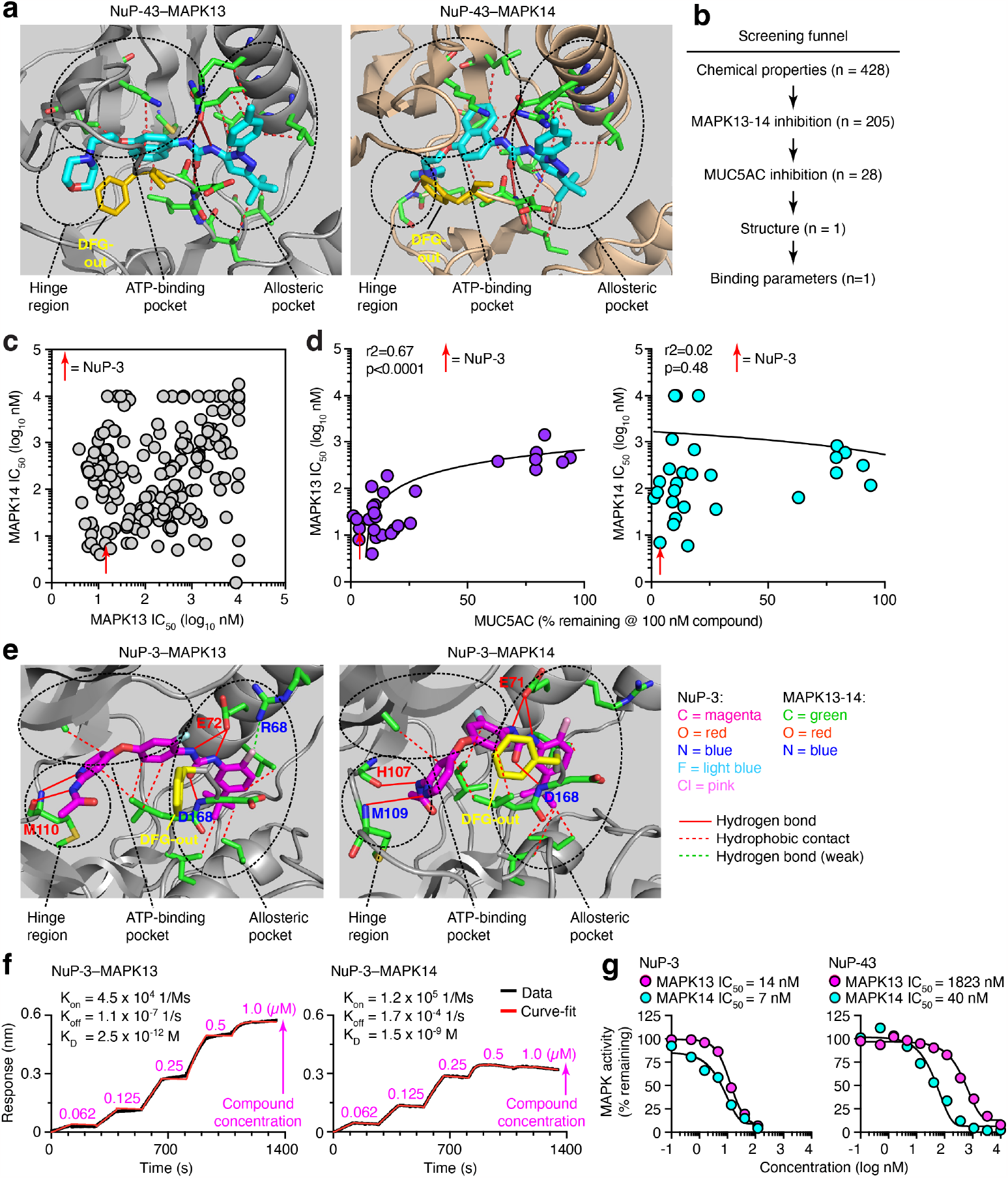
MAPK13-14-guided development of a small-molecule kinase inhibitor. **a**, Structure for parent compound NuP-43 (BIRB-796) docking to MAPK13 (model) and MAPK14 (co-crystal) that illustrates functional targets, hydrogen-bond (solid red lines) and hydrophobic (dashed lines) interactions, and DFG-out binding mode (yellow structure). **b**, Screening funnel of tests for compound library (n=428) to identify candidate compound. **c**, Primary screen of chemically favorable compounds for enzyme IC_50_ for MAPK13 and MAPK14. **d**, Secondary screen for MAPK13 and MAPK14 IC_50_ versus MUC5AC inhibition in IL-13-treated hTECs for selected compounds at 100 nM. **e**, Structure for NuP-3 bound to MAPK13 based on X-ray crystallography and comparable docking to MAPK14 with features labeled as in (**a**) for potential hydrogen-bond interactions for pyridine lone-pair and acetamide-NH with M110/M109/H107 in the hinge region; bidentate urea-NH with E72/E71, urea-O with D168; and chlorine with R68 in the allosteric pocket; and seven sets of hydrophobic contacts. **f**, Biolayer interferometry (BLI) analysis for NuP-3 binding to MAPK13 and MAPK14. **g**, Dose-response for MAPK13 and MAPK14 enzyme inhibition assays for NuP-3.

To better define and predict the function of NuP-3 as a drug candidate, we next obtained the structural basis and corresponding binding kinetics of NuP-3–MAPK13 interactions. Analysis of the X-ray co-crystal structure for the NuP-3–MAPK13 complex revealed a series of hydrogen-bond interactions (pyridine lone-pair and acetamide-NH with M110 in the hinge region; bidentate urea-NH with E72 and urea-O with D168 in the allosteric pocket) (**Fig. 1e** and **Supplemental Table 2**). Comparable interactions are projected for NuP-3 interactions in the hinge region and allosteric pocket for MAPK14 (**Fig. 1e**). In addition, a weak hydrogen bond with R68 in the allosteric pocket in MAPK13 is distinct from the projected interactions with MAPK14 (**Fig. 1e**). However, for both MAPK13 and MAPK14, a lipophilic pocket in the enzyme is filled with the t-butyl-substituted aromatic ring of NuP-3 (**Fig. 1e**). This arrangement requires that the DFG loop of MAPK13 swings out and away from the enzyme (**Fig. 1e**) and thereby stably prevents access to the activation loop. This DFG-out binding mode provides favorable slow dissociation kinetics and consequent high potency and long duration of action of a Type II kinase inhibitor (12, 37, 47). Indeed, bio-layer interferometry (BLI) analysis of NuP-3–MAPK interaction confirmed extremely slow on- and off-rates (hours) for NuP-3 (**Fig. 1f**). This characteristic is similar to NuP-43 (47) binding to MAPK14 that is also based on DFG-out binding mode. Consistent with these findings, enzyme-based assays of NuP-3 activity shows potent inhibition of MAPK13 (IC_50_ = 7 nM) that is increased 130-fold compared to parent compound NuP-43 and retains activity against MAPK14 (IC_50_ = 14 nM) that is in fact increased 6-fold over NuP-43 (**Fig. 1g**).

### NuP-3 effects in human cell models

Based on favorable drug characteristics of NuP-3, further analysis was performed to validate and define utility. To evaluate functional effect, NuP-3 was first tested in hTECs cultured under air-liquid interface conditions with IL-13 stimulation that promotes selective induction of *MAPK13* (versus *MAPK11,12,14,15*) and mucinous differentiation with mucus production marked by expression and secretion of MUC5AC and its mucin granule companion CLCA1 (**Fig. 2a-c**). This protocol requires ALI culture conditions for 21 d to achieve maximal mucus production (**Fig. 2a-c**) and thereby necessitates serial media changes with repeated additions of IL-13 and compound (**Fig. 3a**). Thus, NuP-3 was delivered at 10 nM x 8 doses to achieve significant effects guided by enzyme-based assays (**Fig. 1g**) and was compared to NuP-43 to check for any improvement from the starting chemical scaffold. Under these conditions (that includes increased *MAPK13* expression), NuP-3 treatment markedly inhibited MUC5AC and CLCA1 induction compared to vehicle control and same-dosing with NuP-43 (**Fig. 3b**). These treatments caused no significant effects on the number of viable cells (**Fig. 3c**) consistent with a lack of cell toxicity. We also found a consistent effect of NuP-3 across hTEC cultures established from a series of otherwise healthy subjects (n=7) when tested at 100 nM as a fully effective and still non-toxic concentration (**Fig. 3d,e**). We also tested NuP-3 in 3D cultures when delivered at 10 nM x 5 doses in a protocol optimized for lung organoid formation and mucus production (**Fig. 3f**). Under these conditions, NuP-3 demonstrated similarly effective blockade of IL-13-stimulated *MUC5AC* and *CLCA1* mRNA expression (**Fig. 3g**), again without evidence of cell toxicity based on stable spheroid number and size during treatment (**Fig. 3h**). Together, these results established an effective and nontoxic effect for NuP-3 blockade of mucus production, likely consistent with a role for MAPK13 in controlling mucinous differentiation and mucus production/secretion detected in lung epithelial cell lines and hTECs subjected to shRNA-mediated knockdown of *MAPK13* (but not *MAPK14*) gene expression (12).

**Figure 2.**
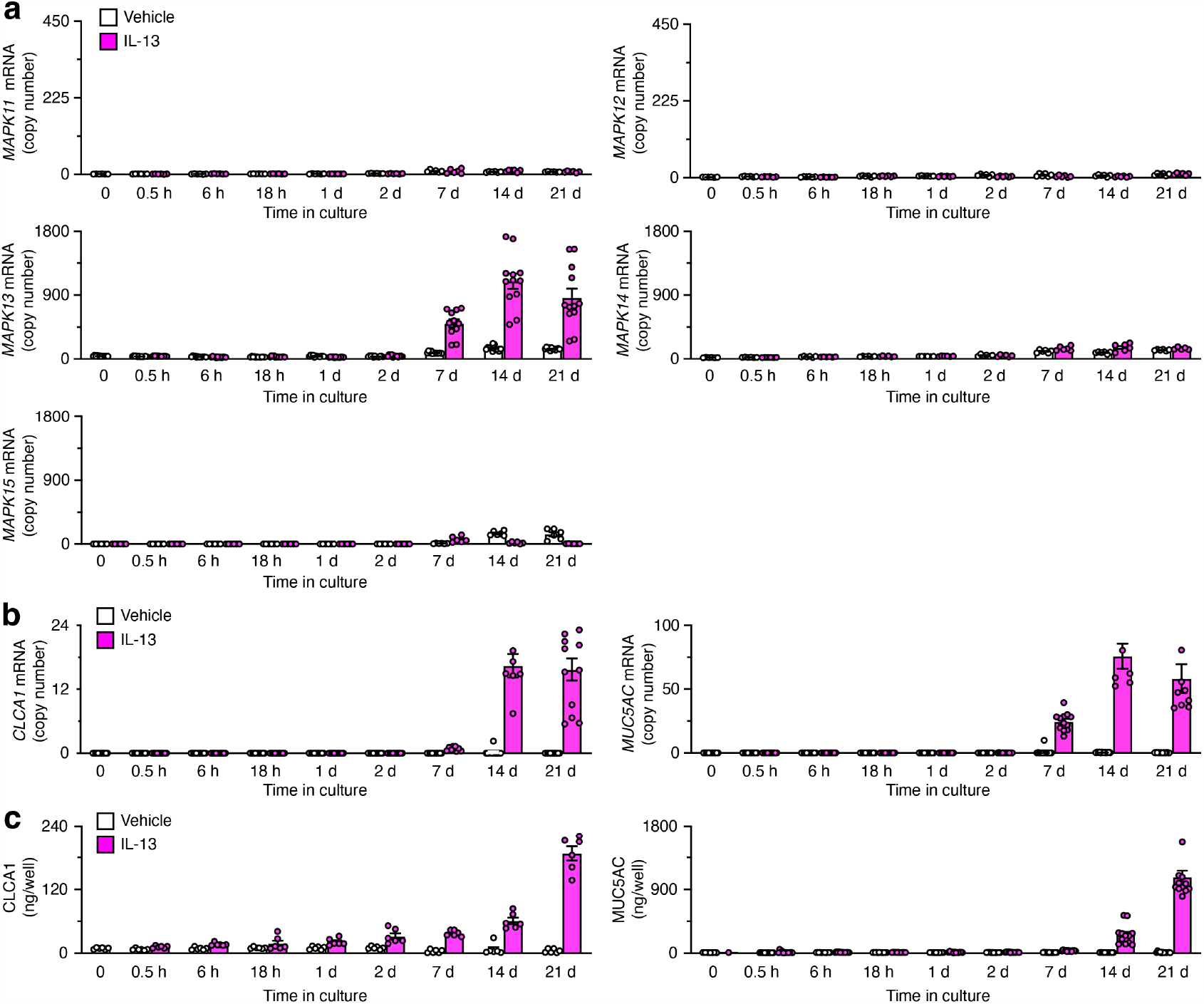
Predominant expression of MAPK13 and correlation with CLCA1 and MUC5AC expression in human airway epithelial cell culture. **a**, Levels of *MAPK11-15* mRNA in human tracheobronchial epithelial cell (hTEC) cultures with and without IL-13 (10 ng/ml) treatment for 0-21 d. **b**, Corresponding level of *CLCA1* and *MUC5AC* mRNA for conditions in (a). **c**, Corresponding levels of CLCA1 and MUC5AC in cell supernatants for conditions in (a). Values were obtained from a single experiment and donor with n=8 samples per condition and are representative of 3 similar experiments. All values represent mean and s.e.m.

**Figure 3.**
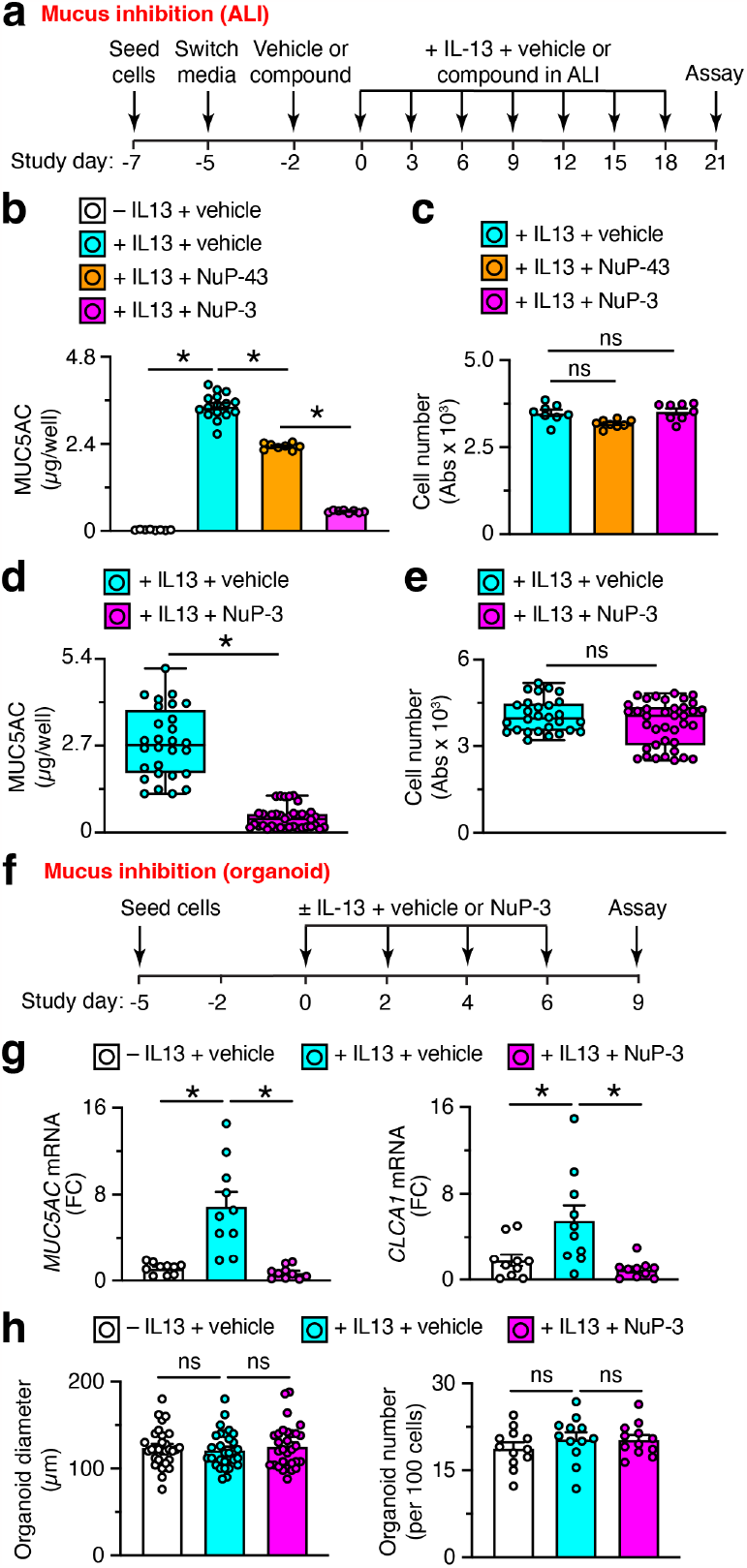
Effect of NuP-3 on mucus production in human airway epithelial cell culture. **a**, Protocol scheme for MUC5AC inhibition for vehicle versus compound: NuP-43 (BIRB-796) or NuP-3 in human tracheobronchial epithelial cells (hTECs) cultured under air-liquid interface (ALI) conditions without or with IL-13 (50 ng/ml). **b**, MUC5AC levels for vehicle versus NuP-43 or NuP-3 (each at 10 nM x 8 doses) for conditions from (a) from one subject representative of three subjects. **c**, Corresponding values for live cell numbers based on resazurin assay for vehicle versus NuP-43 or NuP-3 (each at 100 nM x 9 doses) for conditions in (a) from 1 subject representative of three subjects. **d**, Corresponding values for conditions in (b) derived from 6 subjects. **e**, Corresponding values for conditions in (c) derived from 6 subjects. **f**, Protocol scheme for inhibition of *MUC5AC* and *CLCA1* mRNA for vehicle versus NuP-3 (10 nM x 4 doses) in hTECs cultured under organoid conditions without or with IL-13 (1 ng/ml). FC, fold-change. **g**, Corresponding levels of *MUC5AC* and *CLCA1* mRNA for conditions in (f). Values derived from an individual subject representative of 5 subjects. FC, fold-change. **h**, Corresponding levels of organoid size and number for conditions in (f). Values derived from an individual subject representative of 3 subjects. All bar graphs depict mean ± S.E.M.; box and whisker plots depict upper and lower quartile, range, and median. \**P* <0.05 by ANOVA with Dunnett’s or Tukey correction; ns, not significant. Each sample condition included 4-8 technical replicates per subject.

### NuP-3 efficacy in minipig models

We next advanced NuP-3 to testing in vivo, selecting a minipig model based on relevance to humans given our previous structural and functional genomic comparisons among mouse, pig, and human loci controlling mucus production (48-50). This in vivo approach was also bolstered by PK analysis showing that oral dosing of NuP-3 at 2 mg/kg achieved effective plasma concentrations for at least 8 h in minipigs (**Fig. 4a**). We next needed to establish a pig model of inflammatory lung disease that manifests airway inflammation and mucus production. We arrived at a protocol whereby Yucatan minipigs undergo cytokine challenge using recombinant porcine IL-13 delivered to specific right lung segments (in caudal and accessory lobes) via flexible fiber-optic bronchoscopy on two separate occasions (Study Days 0 and 14) with response monitored in BAL samples (**Fig. 4b**). This approach demonstrated that IL-13-challenge caused reproducible increases in MUC5AC and CLCA1 with no differences between the first and second challenge and therefore no effect of vehicle treatment on the second challenge (**Fig. 4c**, left column). In contrast, these markers of mucus production were significantly decreased by treatment with NuP-3 at 2 mg/kg given orally twice per day for the second challenge (**Fig. 4c**, right column). In addition, treatment with NuP-3 also significantly decreased the influx of immune cells into the airspace (**Fig. 4d**), consistent with blockade of IL-13 induction of chemokine production (12, 33, 36, 51). We also observed attenuation of IL-13-induced MUC5AC, CLCA1, and MUC5B immunostaining in the airway mucosal epithelium without a significant decrease in these same readouts in submucosal glands (**Fig. 5a**). These effects translated to quantitative inhibition of post-challenge MUC5AC, CLCA1, and MUCB staining in mucosal but not submucosal areas using image analysis (**Fig. 5b**). These findings were consistent with CLCA1 control of MUC5AC expression in mucosal epithelium under homeostatic conditions (50).

**Figure 4.**
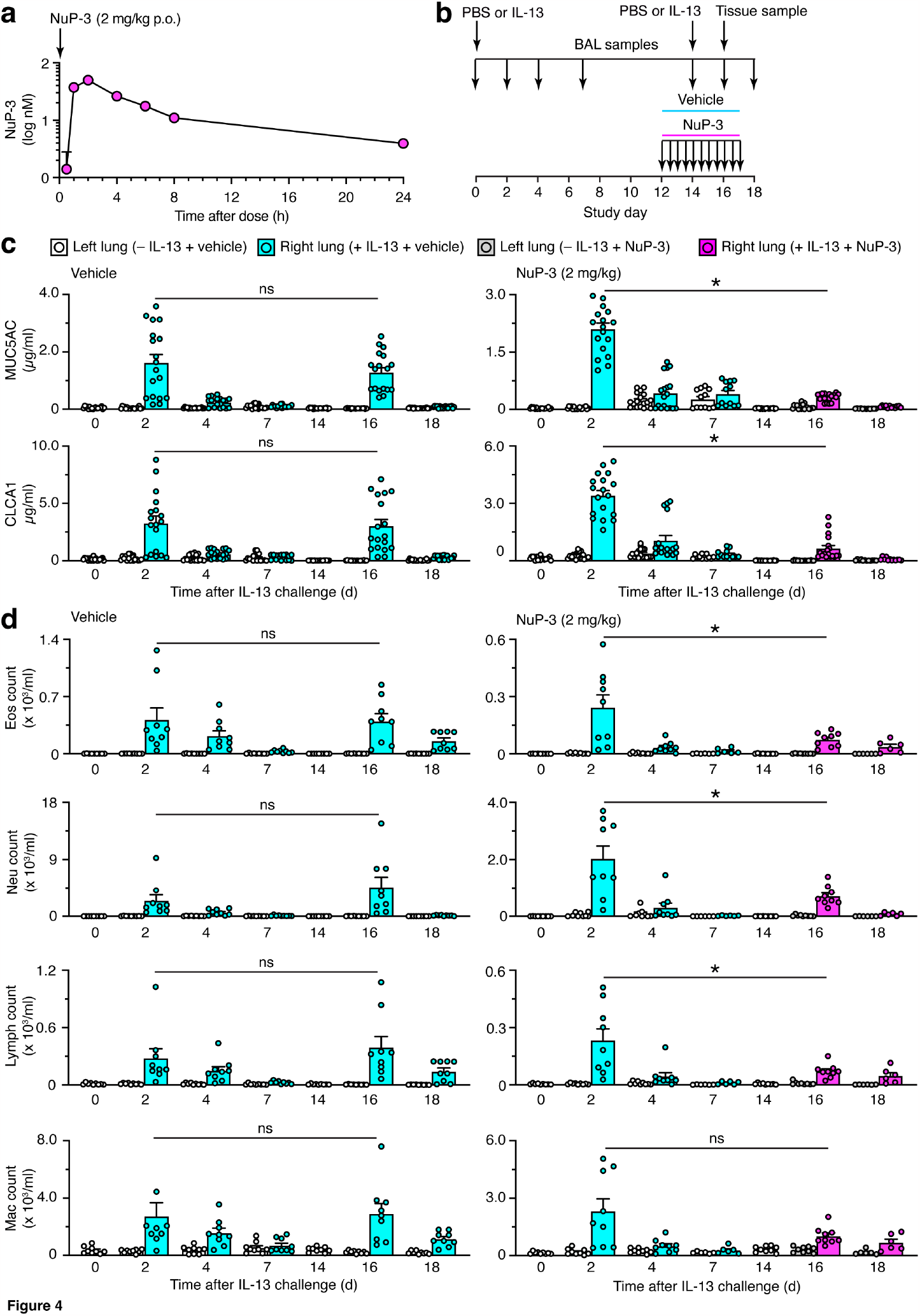
Effect of NuP-3 on IL-13 challenge in minipigs using mucus production and immune cell infiltration readouts. **a**, Pharmacokinetic analysis of NuP-3 in minipigs with plasma concentrations determined after a single oral dose of 2 mg/kg (n=2 minipigs). **b**, Protocol scheme for IL-13 challenge of right lung segments on Study Days 0 and 14 with BAL sample of right and left lung on indicated study days and either vehicle or NuP-3 treatment (2 mg/kg twice per day) on Study Days 12-17. **c**, Levels of MUC5AC and CLCA1 in BAL fluid for protocol scheme in (b) using treatment with vehicle control (left column) or NuP-3 (right column). **d**, Corresponding levels of eosinophil (Eos), neutrophil (Neu), lymphocyte (Lymph), and macrophage (Mac) counts in BAL for conditions in (a). For (c), each plot represents mean ± S.E.M. for 3 minipigs per condition with 3 lung segments and 2 technical replicates per segment per minipig; for (d), values represent 3 minipigs with 3 lung segments and a single assay per segment per minipig. \**P* <0.05 by one-way ANOVA with Tukey correction.

**Figure 5.**
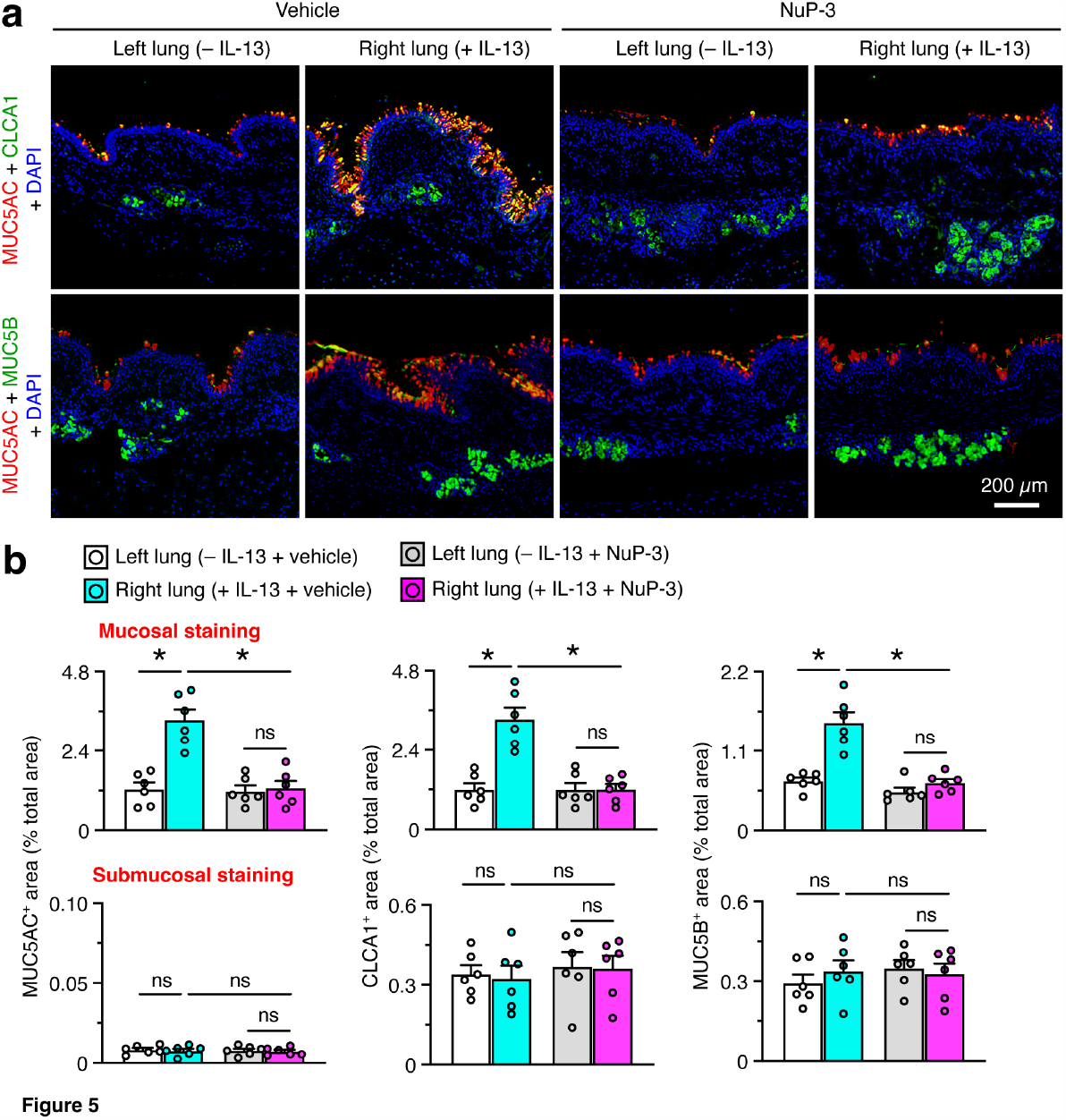
Effect of NuP-3 on lung disease after IL-13 challenge in minipigs using histology readouts. **a**, Immunostaining for MUC5AC plus CLCA1 and MUC5AC plus MUC5B with DAPI counterstaining in lung sections from the protocol scheme in Fig. 3b. Images are representative of 3 minipigs per condition. **b**, Quantitation of mucosal and submucosal immunostaining for conditions in (a). Values represent mean ± S.E.M. for 3 minipigs per condition with 2 technical replicates per minipig. \**P* <0.05 by one-way ANOVA with Tukey correction.

We next engaged the minipig model to determine whether NuP-3 might also block post-viral lung disease. For these experiments, we delivered a natural respiratory pathogen in minipigs (Sendai virus, SeV) (52) to the same right lung segments as for cytokine challenge (**Fig. 6a**). This approach achieved high viral-RNA levels for 2-7 d that were unaffected by treatment with NuP-3 (**Fig. 6b**). In addition, we found that SeV infection caused a significant increase in MUC5AC and CLCA1 levels in BAL samples selectively from the infected site in the right lung, and these increases were markedly attenuated by treatment with NuP-3 at 2 mg/kg given orally twice per day for Study Days -2 to 21 (**Fig. 6c**). Treatment with NuP-3 also significantly inhibited the influx of immune cells into the airspace (**Fig. 6d**), including accumulation of eosinophils as a sign of a type-2 immune response. We also observed attenuation of post-viral increases in MUC5AC, CLCA1, and MUC5B immunostaining in the airway mucosal epithelium without a significant change in these same readouts in submucosal glands (**Fig. 7a**). These effects translated to quantitative blockade of post-viral MUC5AC, CLCA1, and MUC5B staining in mucosal but not submucosal areas (**Fig. 7b**).

**Figure 6.**
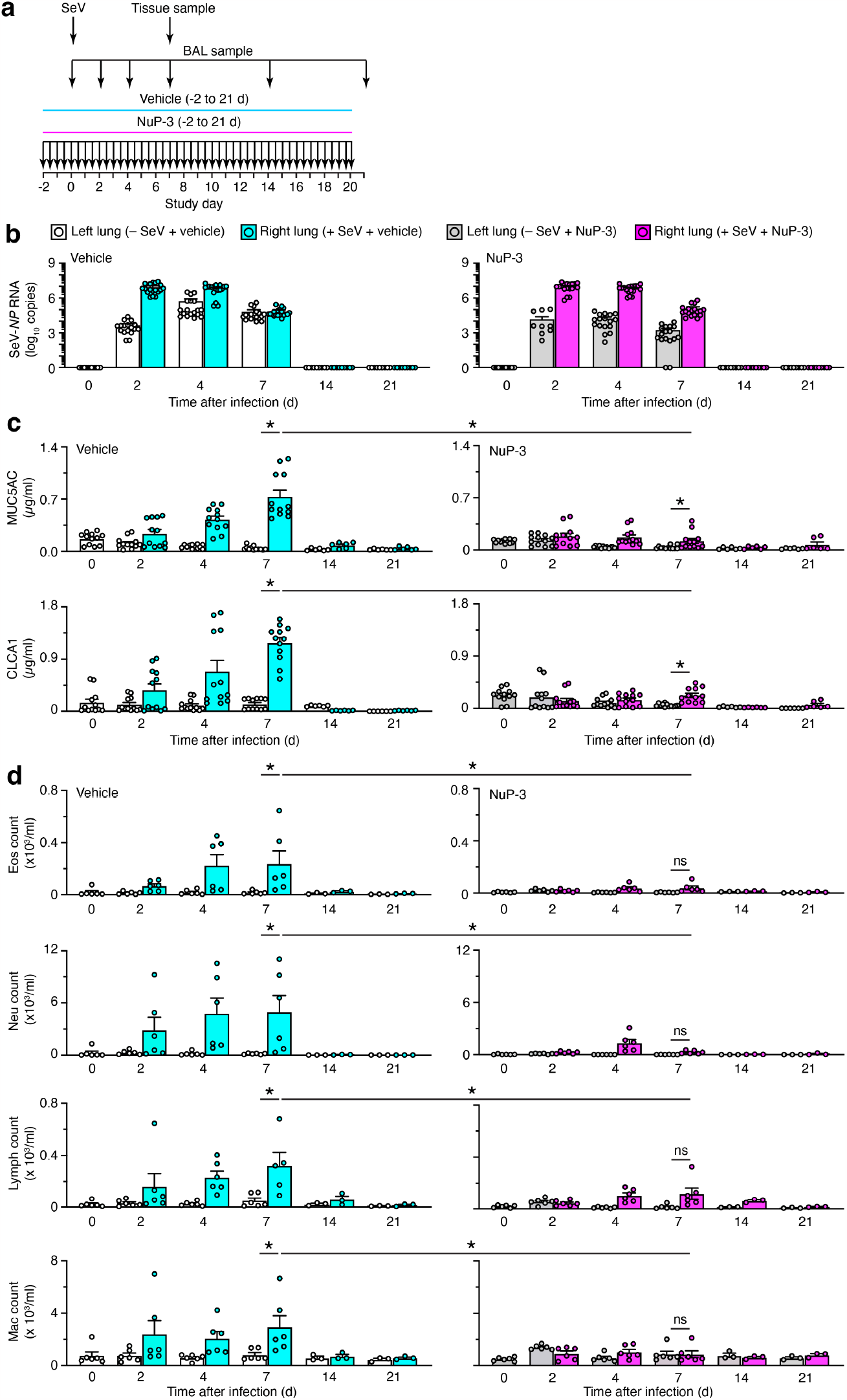
Effect of NuP-3 on lung disease after SeV infection in minipigs using mucus production and immune cell infiltration readouts. **a**, Protocol scheme for SeV delivery into right lung segments on Study Day 0 with tissue and BAL samples of right and left lung on indicated days and either vehicle or NuP-3 treatment at 2 mg/kg orally twice per day on Study Days -2 to 20. **b**, Levels of SeV-*NP* RNA in BAL fluid for scheme in (a) using treatment with vehicle control (left column) or NuP-3 (right column). **c**, Corresponding levels of MUC5AC and CLCA1 in BAL fluid for conditions in (b). **d**, Corresponding levels of eosinophil (Eos), neutrophil (Neu), lymphocyte (Lymph), and macrophage (Mac) counts in BAL for conditions in (b). For (b), each plot represents mean ± S.E.M. for 3 minipigs per condition with 3 lung segments per minipig; for (c), each plot represents 2 minipigs per condition with 3 lung segments and 2 technical replicates per segment per minipig; for (d), values represent 2 minipigs with 3 lung segments and a single assay per minipig. \**P* <0.05 by one-way ANOVA with Tukey correction.

**Figure 7.**
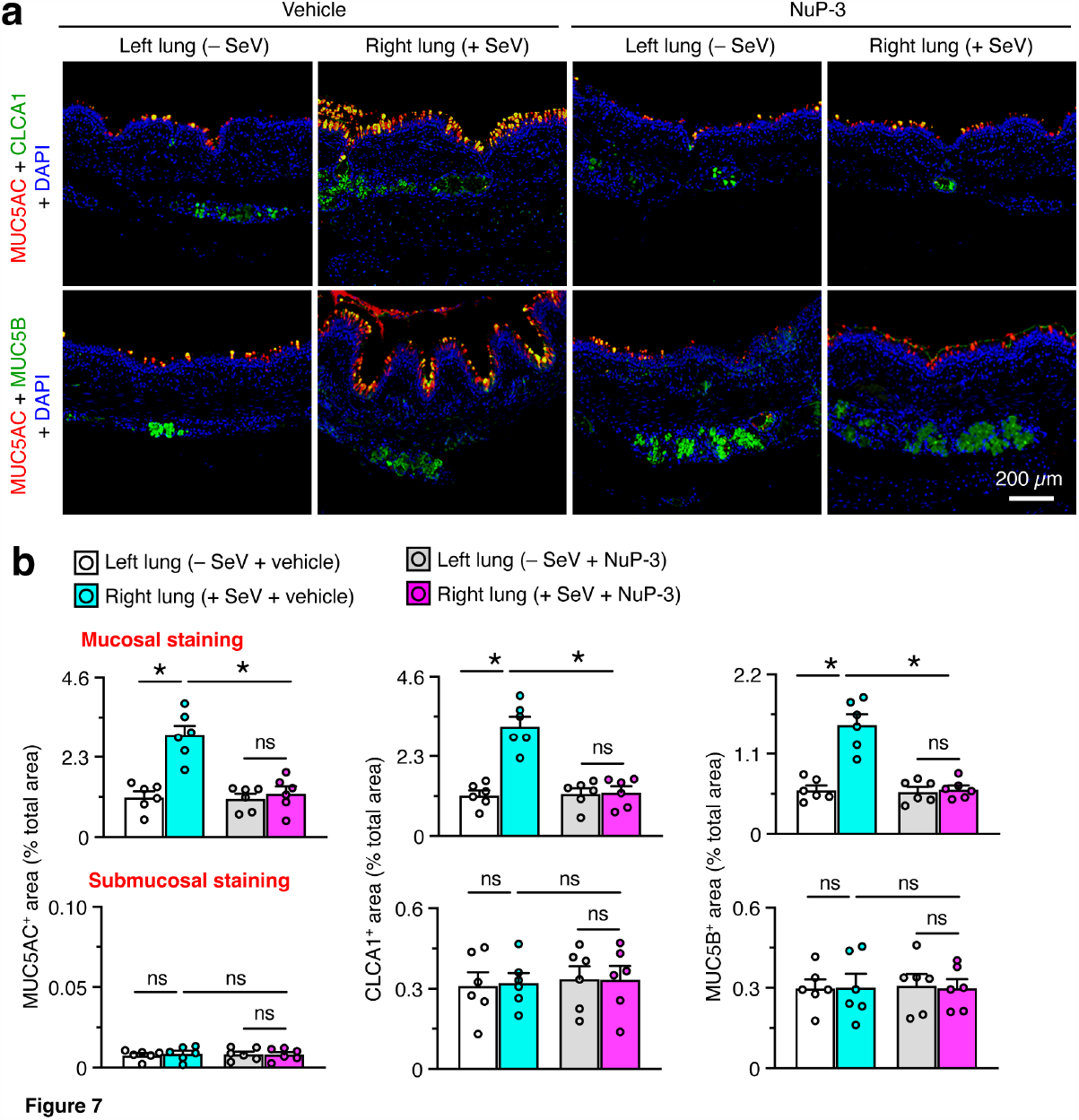
Effect of NuP-3 on lung disease after SeV infection in minipigs using histology readouts. **a**, Immunostaining for MUC5AC plus CLCA1 and MUC5AC plus MUC5B with DAPI counterstaining in lung sections at 7 d after SeV infection from the protocol scheme in Fig. 5a. Images are representative of 3 minipigs per condition. **b**, Quantitation of mucosal and submucosal immunostaining for conditions in (a). Values represent mean ± S.E.M. for 3 minipigs per condition with 2 technical replicates per minipig. \**P* <0.05 by ANOVA with Tukey correction.

To better define mechanism for NuP-3 blockade, we analyzed several biomarkers of disease phenotype in this model. We first observed that there were no significant increases in basal epithelial cells marked by *KRT5* and *AQP3* mRNA levels at the tissue site of infection in the right lung (**Fig. 8a**). This finding is consistent with the short-term timeframe of the minipig model versus the long-term mouse model of post-viral lung disease. In that model, these same biomarkers show Krt5^+^Aqp3^+^ basal-epithelial cell growth is detected at 12 d and is maximal at 21-49 d after SeV infection (33). Nonetheless, the minipig model still showed the same significant increases in *IL13, ARG1*, and *TREM2* mRNA as biomarkers of type 2 inflammation (51, 53), *SERPINB2* and *LTF* mRNA as biomarkers of immune activation of basal epithelial and immune cells (33), and *MUC5AC* and *CLCA1* mRNA as biomarkers of mucinous differentiation (**Fig. 8b-d**). Moreover, each of these increases was significantly down-regulated by treatment with NuP-3 (**Fig 8a-d**), consistent with NuP-3 control of disease-related gene expression. We also detected corresponding increases in SERPINB2 and lactoferrin proteins in BAL fluid, and these increases were also blocked using NuP-3 treatment (**Fig. 8b**). Together, these findings suggested that NuP-3 might act on epithelial and/or immune cells to attenuate airway inflammation and mucus production after respiratory viral infection.

**Figure 8.**
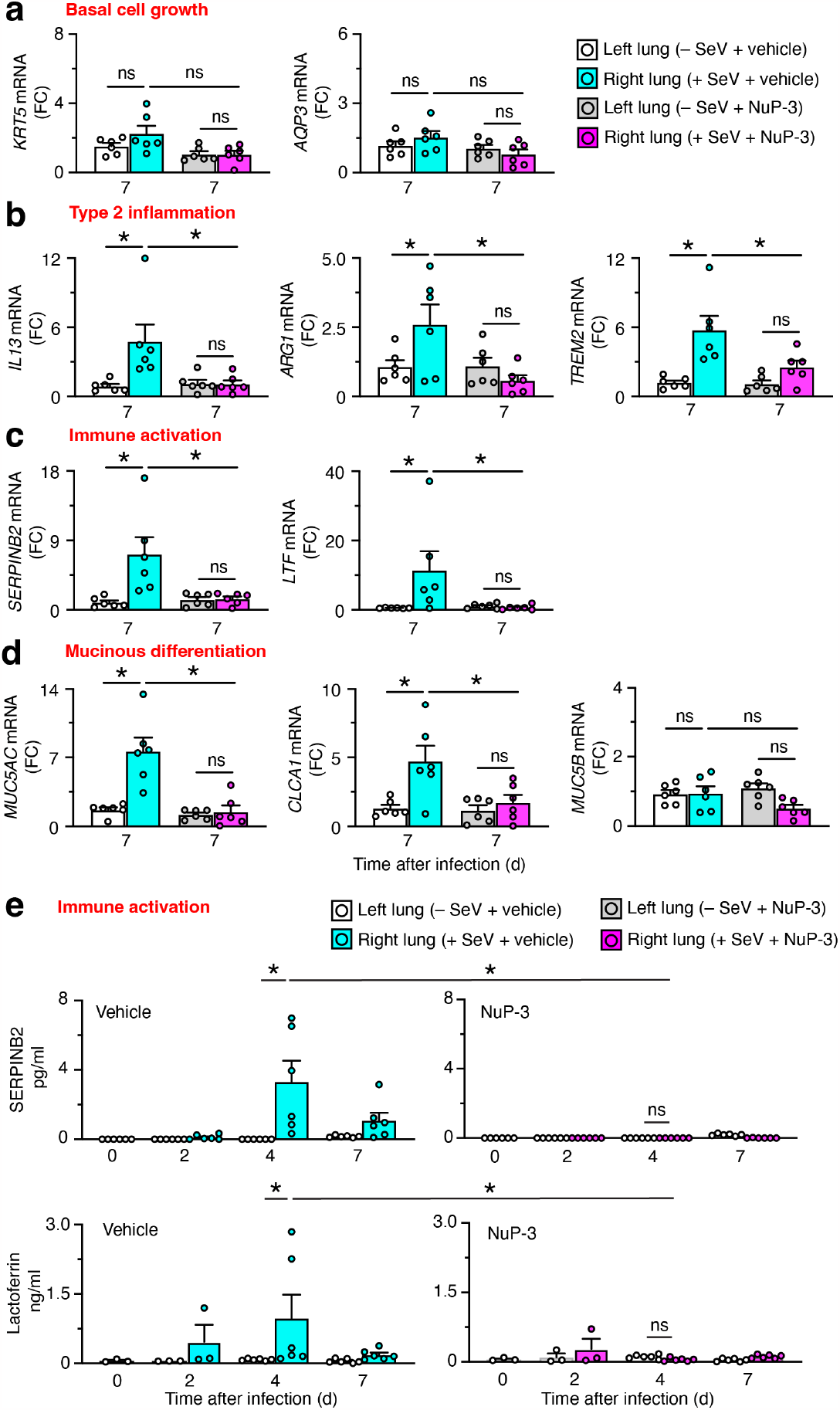
Effect of NuP-3 on lung disease after SeV infection in minipigs using basal epithelial cell-linked readouts. **a-d**, Levels of mRNA biomarkers for basal cell growth (a), type-2 inflammation (b), immune activation (c), and mucinous differentiation (d) in lung tissue at 7 d after SeV infection from the protocol scheme in Fig. 6a. **e**, Corresponding levels of SERPINB2 and lactoferrin proteins in BAL fluid at 0-7 d after SeV infection from the scheme in Fig. 6a. For (a-d), each plot represents mean ± S.E.M. for 3 minipigs per condition with 2 technical replicates per minipig; for (e), values represent 2 minipigs with 3 lung segments and a single assay per segment per minipig. \**P* <0.05 by ANOVA with Tukey correction.

## DISCUSSION

In this study, we use structural biology and structure-based drug-design technologies to target MAPK13-14 activation as a correctable endpoint for respiratory disease. The strategy resulted in the development of a potent MAPK13-14 inhibitor (designated NuP-3) that exhibits favorable drug characteristics and attenuates type-2 cytokine-stimulated mucus production in human airway epithelial cells in culture. In addition, this compound prevents airway inflammation and mucus production in new minipig models driven by type-2 cytokine-challenge (using IL-13) or respiratory viral infection (using SeV). The present data thereby implicate a distinct clinical benefit of a MAPK13-14 inhibitor acting via epithelial and/or immune cells in the airway. To our knowledge, the findings provide the first report of any MAPK13-14 inhibitor that is highly effective in vitro and in vivo. Here we discuss the impact of our findings in five areas related to respiratory disease and inflammatory conditions in general.

The first area concerns the proposed role for MAPK14 as a key component of inflammatory disease and a frequent target for therapeutic intervention based on the canonical role of MAPK14 in inflammatory cytokine signaling and consequent inflammatory disease (14-25). In models of inflammatory disease, MAPK14-specific inhibitors are highly effective in blocking cytokine (e.g., TNF-α and IL-1β) signaling (54). This mechanism predicted therapeutic benefit in a variety of inflammatory conditions including respiratory disease (55); however, even the most advanced versions of these compounds have not proved effective in clinical trials of patients with lung disease, e.g., COPD (18). The data here suggest a basis for this failure by implicating the additional action of MAPK13 in the disease process. As predicted from our earlier work (12), MAPK13 activation again appears required for type-2 cytokine (particularly IL-13 and IL-4) signaling as a key driver of inflammation and mucus production. Our models were not designed to specifically test MAPK14-dependent events, but blockade of inflammatory cytokine signaling remains a target for immune-based lung disease, including Covid-19 (56). In that context, we recognize that NuP-3 maintains potent MAPK14-blocking activity. Indeed, this compound was designed to gain MAPK13 and not yet eliminate MAPK14 inhibition. Thus, the present data suggest that attacking MAPK13-14 together might achieve an unprecedented therapeutic benefit for the host response to a broader range of cytokine-signaling and viral-infection conditions. However, NuP-3 was more effective than NuP-43 despite similar potencies for MAPK14 inhibition, and our earlier MAPK gene knockdown work showed no significant effect of MAPK14 blockade on mucus production (12). Additional studies of targeted gene knockout and highly selective inhibitors will be needed to fully define MAPK13 function alone and in combination with MAPK14.

In that regard, a second area of impact for the present data relates to role of the type-2 immune response for host defense and inflammatory disease. In our previous work, we identified basal-ESC growth, immune activation, and mucinous differentiation as requirements for long-term lung remodeling disease after SeV infection in mice (33). In that case, however, we observed basal-ESC hyperplasia and metaplasia at bronchiolar-alveolar sites in the setting of a more severe and widespread infection. Here we find the same markers of basal-epithelial cell activation (SERPINB2 and lactoferrin), but lung disease is limited to an airway site for mucinous differentiation as part of a milder and likely more localized infection. Moreover, these basal-epithelial cell markers are shared with immune cells, so the target site(s) for blocking immune activation still need to be defined. Additional work is also needed to define whether more severe viral infection can drive more pronounced remodeling disease in large animals and humans, but this already seems to be the case in Covid-19 patients (36).

Relevant to this point, a third area of interest relates to the timing of events in the development of inflammatory disease. In that regard, respiratory viral infection often progresses as a top-down infection that starts in the upper and lower airways and extends distally to bronchiolar-alveolar sites. Similarly, acute inflammation due to general immune activation (including TNF-α, IL-1β, and type 1 and type 2 cytokine actions) might be followed by progressive and prolonged type 2 cytokine production. This paradigm is consistent with our observations in mouse models and Covid-19 patients (36), but here again further work will be needed to establish this pattern and define functional consequences. The present data suggests that combined blockade that includes MAPK13 inhibition is of unexpected benefit even for relatively short-term airway inflammation and mucus production and thereby adds to our earlier work in long-term lung remodeling disease.

Related to this issue, a fourth area of impact is the relationship of the present findings to the sequential process of drug discovery and development. Thus, the present work moves significantly closer to clinical application by increasing the potency of MAPK13 inhibitors beyond previous reports (12, 57). The present success versus previous screening attempts derives from optimization work using a NuP-43–MAPK14 co-crystal structure to create a model of NuP-43–MAPK13 and thereby aid the design of improved MAPK13 inhibitors. Then, at the conclusion of this work, our own NuP-3–MAPK13 co-crystal structure was used to define the basis for optimized compound binding into MAPK13. From this information, we could create an analogous model of NuP-3– MAPK14 to also explain the binding interactions for this complex. This process thereby resulted in potent biochemical and biologic activities for NuP-3. However, further preclinical studies are needed ahead of clinical trials in humans. This data includes additional pharmacology to establish optimal dosing level, route, and timing in relation to disease prevention and reversal. In addition, safety pharmacology and toxicology studies in preclinical species will be needed to establish therapeutic index and adverse effect level. In concert with this process, the ongoing analysis of newly identified MAPK13-inhibitor interactions will guide further development of the MAPK13-targeted drug pipeline.

A fifth and final area of interest relates to the practical need for more successful therapy of post-viral lung disease and related inflammatory conditions. Respiratory viruses are perhaps the most common cause of medical attention for infectious illness, particularly during pandemic conditions such as the outbreaks of influenza virus or coronavirus infections (58, 59). Further, even after clearance of infectious virus, the acute illness can progress to respiratory failure in the intermediate term and to chronic respiratory disease in the long term (34, 60). This paradigm is also relevant to virus-triggered disease in asthma, COPD, and Covid-19 (12, 13, 36, 51, 61-63). Despite the magnitude of these public health problems, there is still a need for a precisely designed small-molecule drug to attenuate these disease conditions. The present work suggests a method to modify the epithelial and/or immune cell populations linked to short-term airway disease manifest by excessive inflammation and mucus production (33, 53, 64, 65). Thus, we identify a more potent MAPK13-14 inhibitor that achieves a remarkable benefit for the host response to type 2 cytokine-challenge or respiratory viral infection as a model for this type of disease. Additional study will be needed to address whether the same therapeutic approach will modify long-term disease that includes basal-ESC reprogramming and immune-cell activation (33, 53, 64, 65). Nonetheless, the present insight provides the next useful step in developing a safe and effective drug for respiratory disease and other diseases that feature increases in MAPK13 and/or MAPK14 expression and activation (66).

## Abbreviations used in this article

BAL: bronchoalveolar lavage
BLI: biolayer interferometry
basal-ESC: basal-epithelial stem cell
CLCA1: chloride channel accessory 1
COPD: chronic obstructive pulmonary disease
Covid-19: coronavirus disease of 2019
hTEC: human tracheobronchial epithelial cell
MAPK: mitogen-activated protein kinase
MUC5AC: mucin 5AC
MUC5B: mucin 5B
SeV: Sendai virus.

## DATA AVAILABILITY

The data that support the findings of this study will be made available upon reasonable request to the corresponding author.

## SUPPLEMENTAL DATA

Supplemental Tables 1-2: https://doi.org/10.6084/m9.figshare.23464478.

Private link: https://figshare.com/s/42f4a4985f372caf4918

## ACKNOWLEDGMENTS

The authors thank the Pulmonary Morphology Core, Division of Comparative Medicine, and Eugene Agapov, Yael Alevy, Rowena Grainger, Olga Korkovsky, and Michael Talcott for research support.

## AUTHOR CONTRIBUTIONS

SPK, KW, YZ, DM, ML, CAI, SRA, JY, SP, SLB, JRC, ZH, DEB, and AGR performed experiments and analyzed data; MJH designed the project, analyzed data, and wrote the manuscript; and all authors approved the final version of the manuscript.

