## Supplemental Data for "A potent MAPK13-14 inhibitor prevents airway inflammation and mucus production"

**Supplemental Table 1**. Sequences of DNA primers and probes for real-time qPCR assays in human and mini-pig samples.

| **Target Gene** | **Species** | **Type** | **ID/Sequence** |
| --- | --- | --- | --- |
| *GAPDH* | Human | F  R  P | 5’-CAGCCGAGCCACATCCCTCAGACACCAT-3’  5’-CTTTACCAGAGTTAAAAGCAGCCCTGGTGACCA-3’  5’-AGGTCGGAGTCAACCGATTTGGTCGTATTG-3’ |
| *MUC5AC* | Human | F  R  P | 5’-AGGCCAGCTACCGGGCCGGCCAGACCAT-3’  5’-GTCCCCGTACACGGCGCAGGTGGCCAGGCA-3’  5’-TGCAACACCTGCACCTGTGACAGCAGGAT-3’ |
| *ARG1* | Mini-pig |  | Ss03391394_m1 (ThermoFisher Scientific) |
| *CLCA1* | Mini-pig | F  R  P | 5’-CTGACGTGGACGGCTCCTGGGGAT-3’  5’-GAACTTGTCTCTGAGATCAAGAATATTTGTGCT-3’  5’-TTACGACCACGGAAGAGCTGACAGGT-3’ |
| *GAPDH* | Mini-pig | F  R  P | 5’-CCCCGCGATCTAATGTTCT-3’  5’-CTTCACCATCGTGTCTCAGG-3’  Probe #6 (Roche Universal Probe Library) |
| *IL13* | Mini-pig | F  R  P | 5’-TGCTGAGCGCCCTCTGTTCTCACAAGC-3’  5’-TACGAACTGGGCCACTTCAATTTTGGTG-3’  5’-CAAGCGAGCAAGTTCCTGGCAAGCACATCCGA-3’ |
| *LTF* | Mini-pig |  | Ss03384354_u1 (ThermoFisher Scientific) |
| *MUC5AC* | Mini-pig | F  R  P | 5’-CGTAGAGCACAGGTGCAAGT-3’  5’-GCAGGGTCACGTTTCTCAG-3’  Probe #17 (Roche Universal Probe Library) |
| *MUC5B* | Mini-pig | F  R  P | 5’- cagccaacaactgcacaga -3’  5’- cagccaacaactgcacaga -3’  Probe #3 (Roche Universal Probe Library) |
| *SERPINB2* | Mini-pig |  | Ss06916366_m1 (ThermoFisher Scientific) |
| *TREM2* | Mini-pig | F  R  P | 5’- ctagtagccagcgccctct -3’  5’- tggggaagtcctctgtttgt -3’  Probe #64 (Roche Universal Probe Library) |
| *SeV-NP* | Not applicable | F  R  P | 5’-GGCGGTGGTGCAATTGAG-3’  5’-CATGAGCTTCTGTTTCTAGGTCGAT-3’  5’-AGCTCTAGACAATGCC-3’ |

^1^Abbreviations: F, forward primer; R, reverse primer; P, MGB probe.

**Supplemental Table 2.** Data collection and refinement statistics for the NuP-3–MAPK13 co-crystal complex.

| **Statistic** | **Value** |
| --- | --- |
| **Data collection** | |
| Space group | P2_1_ |
| Cell dimensions |  |
| *a, b, c* (Å) | 75.5, 83.4, 134.5, 98.3 |
| Resolution (Å) | 46.48-3.15(3.26-3.15) |
| R_merge_ |  |
| *I / σ* | 4.9(0.8) |
| *CC_1/2_* | 0.99(0.36) |
| Completeness (%) | 93.9(95.0) |
| Redundancy | 3.7(3.5) |
| **Refinement** | |
| Resolution (Å) | 46.5~3.15 |
| No. reflections | 23873(2770) |
| *R*_work_ / *R*_free_ | 0.264(0.364)/0.324(0.416) |
| No. atoms | 10842 |
| Protein | 10710 |
| Ligand/ion | 132 |
| Water | N/A |
| *B*-factors | 124.7 |
| Protein | 125.1 |
| Ligand/ion | 95.0 |
| Water |  |
| R.m.s. deviations |  |
| Bond lengths (Å) | 0.005 |
| Bond angles (°) | 1.13 |
